# SIRT2, ERK and Nrf2 mediate NAD+ treatment-induced increase in the antioxidant capacity of differentiated PC12 cells under basal conditions

**DOI:** 10.1101/497404

**Authors:** Jie Zhang, Yunyi Hong, Wei Cao, Haibo Shi, Weihai Ying

**Affiliations:** Shanghai Sixth People’s Hospital and School of Biomedical Engineering, Shanghai Jiao Tong University, Shanghai, P.R. China; Department of Otorhinolaryngology, Shanghai Sixth People’s Hospital Affiliated to Shanghai Jiao Tong University, Shanghai, P.R. China

**Keywords:** NAD^+^, GSH, Nrf2, SIRT2, ERK

## Abstract

NAD^+^ administration is highly beneficial in numerous models of diseases and aging. It is becoming increasingly important to determine if NAD^+^ treatment may increase directly the antioxidant capacity of cells under basal conditions. In current study we tested our hypothesis that NAD^+^ can enhance directly antioxidant capacity of cells under basal conditions by using differentiated PC12 cells as a cellular model. We found that NAD^+^ treatment can increase the GSH/GSSG ratios in the cells under basal conditions. NAD^+^ can also increase both the mRNA and protein level of γ-glutamylcysteine ligase – a key enzyme for glutathione synthesis, which appears to be mediated by the NAD^+^-induced increase in Nrf2 activity. These NAD^+^-induced changes can be prevented by both SIRT2 siRNA and the SIRT2 inhibitor AGK2. The NAD^+^-induced changes can also be blocked by the ERK signaling inhibitor U0126. Moreover, the NAD^+^-induced ERK activation can be blocked by both SIRT2 siRNA and AGK2. Collectively, our study has provided the first evidence that NAD^+^ can enhance directly the antioxidant capacity of the cells under basal conditions, which is mediated by SIRT2, ERK, and Nrf2. These findings have suggested not only the great nutritional potential of NAD^+^, but also a novel mechanism underlying the protective effects of the NAD^+^ administration in the disease models: The NAD^+^ administration can enhance the resistance of the normal cells to oxidative insults by increasing the antioxidant capacity of the cells.

## Introduction

NAD^+^ plays critical roles in energy metabolism, mitochondrial functions, calcium homeostasis and immune functions (Ying, 2005; Ying, 2008; Ma et al., 2012). A number of studies have shown that NAD^+^ administration is highly beneficial in multiple models of major diseases (Ma et al., 2012). Moreover, NAD^+^ and its precursors such as nicotinamide mononucleotide (NMN) can produce significant anti-aging effects (Scheibye-Knudsen et al., 2014; Fang et al., 2016). Cumulating evidence has suggested that NAD^+^ deficiency is a common pathological factor of a number of diseases and aging (Ma et al., 2012). It becomes increasingly important to further elucidate the mechanisms underlying the beneficial effects of NAD^+^ administration in diseases and aging, which is necessary for not only elucidation of the roles of NAD^+^ administration in diseases and aging, but also clinical and nutritional applications of NAD^+^.

Oxidative stress plays critical pathological roles in numerous diseases and aging (Halliwell, 2005; Lin and Beal, 2006). Multiple studies have shown that NAD^+^ treatment can decrease oxidative stress-induced death of several cell types including neurons, astrocytes and myocytes (Ying et al., 2003; Alano et al., 2004; Alano et al., 2010; Zhang et al., 2016). Several studies have suggested that NAD^+^ treatment can indirectly decrease oxidative cell death by such mechanisms as enhancement of SIRT1 activity and prevention of glycolytic inhibition, mitochondrial permeability transition, mitochondrial depolarization and nuclear translocation of apoptosis-inducing factor (AIF) (Alano et al., 2004; Araki et al., 2004; Alano et al., 2010; Hong et al., 2014).

We proposed a hypothesis regarding the new mechanism underlying the protective effects of NAD^+^ administration in the disease models: NAD^+^ may enhance the antioxidation capacity of normal cells, which would increase the resistance of the cells to oxidative insults in the disease models. In our current study, we tested our hypothesis that NAD^+^ can enhance directly antioxidant capacity of cells under basal conditions by using differentiated PC12 cells as a cellular model. Demonstration of this hypothesis would also suggest valuable information indicating the nutritional potential of NAD^+^. Our study has indicated that NAD^+^ can directly increase the GSH/GSSG ratio of differentiated PC12 cells by a γ-glutamylcysteine ligase (GCL)-dependent mechanism, which is mediated by SIRT2, ERK, and Nrf2.

## Materials and Methods

### Cell cultures

Cell cultures were prepared as described previously (Chen et al., 2013). Differentiated PC12 cells were purchased from the Cell Resource Center of Shanghai Institute of Biological Sciences, Chinese Academy of Sciences. The cells were plated onto 6-well or 24-well cell culture plates at the initial density of 1×10^5^ cells/ml in Dulbecco’s Modified Eagle Medium (DMEM) containing 4500 mg/L D-glucose, 584 mg/L L-glutamine, 110 mg/L sodium pyruvate (Thermo Scientific, Waltham, MA, USA), which also contains 1% penicillin and streptomycin (Invitrogen, Carlsbad, CA, USA) and 10% fetal bovine serum (PAA, Linz, Austria). The cells were used when the density of the cell cultures reached 60-80%.

### Experimental procedures

Experiments were initiated by replacing the cell culture medium with medium containing various concentrations of drugs. The cells were left in an incubator with 5% CO_2_ at 37 °C for various durations.

### SIRT2 RNA silencing

The density of differentiated PC12 cells was approximately 50% at the time of transfection. Three small interfering RNA (siRNA) duplexes against rat SIRT2 (NM_001008368) at nucleotides 754 – 772 (CC UUGCUAAGGAGCUCUAUTT), 843 – 862 (GCUGC UACACGCAGA AUAUTT), and 1393 – 1411 (GGAGCAUGCCAACAUAGAUTT) were commercially synthesized (Genepharma, Shanghai, China). Scrambled RNA oligonucleotides were used for control cells. The transfection was conducted according to the manufacturer’s instructions. For each well, 600 μl serum-free media containing 33.33 nM of each of the three SIRT2 siRNA oligos and 4 μl lipofectamine 2000 (Invitrogen, Carlsbad, CA, USA) were added. After incubation for 6 hours, the media was replaced by DMEM containing 10% fetal bovine serum.

### DHE assay

Dihydroethidium (DHE, Beyotime Institute of Biotechnology, Jiangsu, China) assay were used to determine the reactive oxidative stress. Cells were treated for 6 hours, then harvested, washed with serum-free DMEM and incubated with 10 μmol/l DHE for 20 minutes at 37 °C. Then cells were fixed with 4% paraformaldehyde for 10 minutes and stained with DAPI (1:1000 dilution, Beyotime Institute of Biotechnology, Jiangsu, China). Subsequently, the cells were washed with PBS, and the fluorescence signals of the cells were observed under a Leica fluorescence microscope.

### Determinations of GSH, GSSG, and GSH/GSSG ratios

The levels of total glutathione (GSH+GSSG) and GSSG were determined by using a commercially available kit (Beyotime Institute of Biotechnology, Haimen, China), which was conducted according to the manufacturer’s instructions. In brief, cells were washed with PBS for three times and lysed, then frozen at −80 °C and thawed at room temperature for three times. The levels of total glutathione were assessed by the 5, 5’ dithiobis (2-nitrobenzoic acid) (DTNB)-oxidized glutathione (GSSG) reductase-recycling assay. In brief, 700 μl of 0.33 mg/ml NADPH in 0.2 M sodium phosphate buffer containing 10 mM ethylenediaminetetraacetic acid (pH 7.2), 100 μl of 6 mM DTNB, and 190 μl of distilled water were added and mixed in an eppendoff tube. The reaction was initiated by addition of 10 μl of 250 IU/ml glutathione reductase. With readings recorded every 30 seconds, the absorbance was monitored at 412 nm by a plate reader for 15 minutes. To determine the levels of GSSG, 10 μl samples were vigorously mixed with 0.2 μl of 2-vinylpyridine and 0.6 μl triethanolamine. After one hour, the sample was assayed as described above in the DTNB-GSSG reductase-recycling assay. According to the methods described above, standards (0.01, 0.02, 0.1, 0.2, and 1.0 mM) of total glutathione or GSSG were also assessed. The amount of total glutathione and GSSG in each sample was normalized by the protein concentration of the sample that was determined by BCA Protein Assay (Thermo Scientific, Waltham, MA, USA). The GSH concentrations were calculated from the differences between the concentrations of total glutathione and the concentrations of GSSG.

### Western blots

Western blot assays were conducted according to a protocol reported in a previous study (Nie et al., 2014). Cells were lysed in RIPA buffer (Millipore, Temecula, CA, USA) containing Complete Protease Inhibitor Cocktail (Roche Diagnostics, Mannheim, Germany) as well as 1 mM PMSF, 30 μg of total protein was loaded for each sample, which were electrophoresed through a 10% SDS-polyacrylamide gel, and then transferred to 0.45 μm nitrocellulose membranes (Millipore, CA, USA) on a semi-dry electro transferring unit (Bio-Rad Laboratories, CA, USA). Membranes were blocked in 5% bovine serum albumin (BSA) in TBST for 2 hours at room temperature and then incubated with GCLc antibody (1:1000 dilution, 12601-1-AP; Proteintech Group, Chicago, Illinois, USA), GCLm antibody (1:1000 dilution, 14241-1-AP; ProteinTech Group, Chicago, Illinois, USA), or Nrf2 antibody (1:1000, C-20; Santa Cruz Biotechnology, Heidelberg, Germany), p-ERK antibody (1:2000 dilution, 4370; CST, Boston, MA, USA), ERK antibody (1:2000 dilution, 4695; CST), SIRT2 antibody (1:1000 dilution, sc-20966; Santa Cruz Biotechnology, Dallas, TX, USA), β-tubulin (1: 2000 dilution, M20005; Abmart, Shanghai, China) or Lamin A/C (1: 1000 dilution, 2032; CST) overnight at 4 °C. Subsequently, the membranes were incubated with appropriate HRP-conjugated secondary antibody (Epitomics, Hangzhou, Zhejiang Province, China). The protein signals were visualized by using an ECL detection system (Pierce Biotechnology, Rockford, IL, USA). β-tubulin levels were used to normalize sample loading and transfer and Lamin A/C levels were used to normalize nuclear protein samples. Gel-Pro Analyzer was used to quantify the bands.

### Quantitative real-time PCR assays of the mRNA levels of GCLc, GCLm and Nrf2

Quantitative real-time PCR assays were performed as described previously (Ma et al., 2011; Xie et al., 2013). After total RNA was extracted (TaKaRa MiniBEST Universal RNA Extraction Kit, Takara Bio, Dalian, China) from PC12 cells, 500 ng total RNA was reverse-transcribed to cDNA (Prime-Script RT reagent kit, Takara Bio, Dalian, China). RT-PCR was performed according to the following procedure: 37 °C for 15 minutes, then 85 °C for 15 seconds. Quantitative real-time PCR was performed by using SYBR Premix Ex Taq (Takara Bio, Dalian, China) and the following primers: Nrf2 (sense 5’-GACCTAAAGCACAGCCAACACAT-3’ and anti-sense 5’-CTCAATCGGCTTGAATGTTTGTC-3’); GCLc (sense 5’-ATGGAGGTACAGTTGACAG AC-3’ and anti-sense 5’-ACGGCGTTGCCACCTTTGCA-3’); GCLm (sense 5’-GCTGTACCAGT GGGCACAG-3’ and anti-sense 5’-GGCTTCAATGTCAGGGATGC-3’); GAPDH (sense 5’-AGGTCGGTGTGAACGGATTTG-3’ and anti-sense 5’-TGTAGACCATGTAGTTGA GGTCA-3’). PCRs were performed according to the following procedure: After denatured at 95 °C for 10 s, 40 cycles of 95 °C for 5 s and 60 °C for 30 s. The data was analyzed by using the comparative threshold cycle method, and results were expressed as fold difference normalized to the level of GAPDH mRNA.

### Determinations of intracellular NAD^+^ concentrations

As previously described (Ying et al., 2003), NAD^+^ concentrations were determined by the recycling assay. Briefly, samples were extracted in 0.5 N perchloric acid. After centrifugation at 12,000 RPM for 5 minutes, the supernatant was obtained, which was neutralized to pH 7.2 by using 3 N potassium hydroxide and 1 M potassium phosphate buffer. After centrifugation at 12,000 RPM for 5 minutes, the supernatants were mixed with a reaction media containing 1.7 mg 3-[4,5-dimethylthiazol-2-yl]-2,5-diphenyl-tetrazolium bromide (MTT), 1.3 mg alcohol dehydrogenase, 488.4 mg nicotinamide, 10.6 mg phenazine methosulfate, and 2.4 mL ethanol in 37.6 mL Gly-Gly buffer (65 mM, pH 7.4). After 10 minutes, the A_560nm_ was determined by a plate reader, and the readings were calibrated with NAD^+^ standards. The protein concentrations were assessed by BCA Protein Assay (Thermo Scientific, Waltham, MA, USA).

### Statistical analyses

All data are presented as mean ± SE. Data were assessed by one-way ANOVA, followed by Student-Newman-Keuls *post hoc* test. *P* values less than 0.05 were considered statistically significant.

## Results

GSH/GSSG ratio is a major index of antioxidant capacity of cells (Dickinson and Forman, 2002). Our study determined the effects of NAD^+^ on the GSH/GSSG ratio and the levels of GSH, GSSG and total glutathione (GSH + GSSG) in the PC12 cells under basal conditions: Treatment of the cells with 0.01, 0.1 and 1 mM NAD^+^ dose-dependently increased the GSH levels of the cells (Fig. 1A), while only 1 mM NAD^+^ was capable of mildly increased the GSSG levels (Fig. 1B). One mM NAD^+^ significantly increased the GSH/GSSG ratio (Fig. 1C), while both 0.1 and 1 mM NAD^+^ significantly increased the total glutathione levels (Fig. 1D).

**Figure 1.**
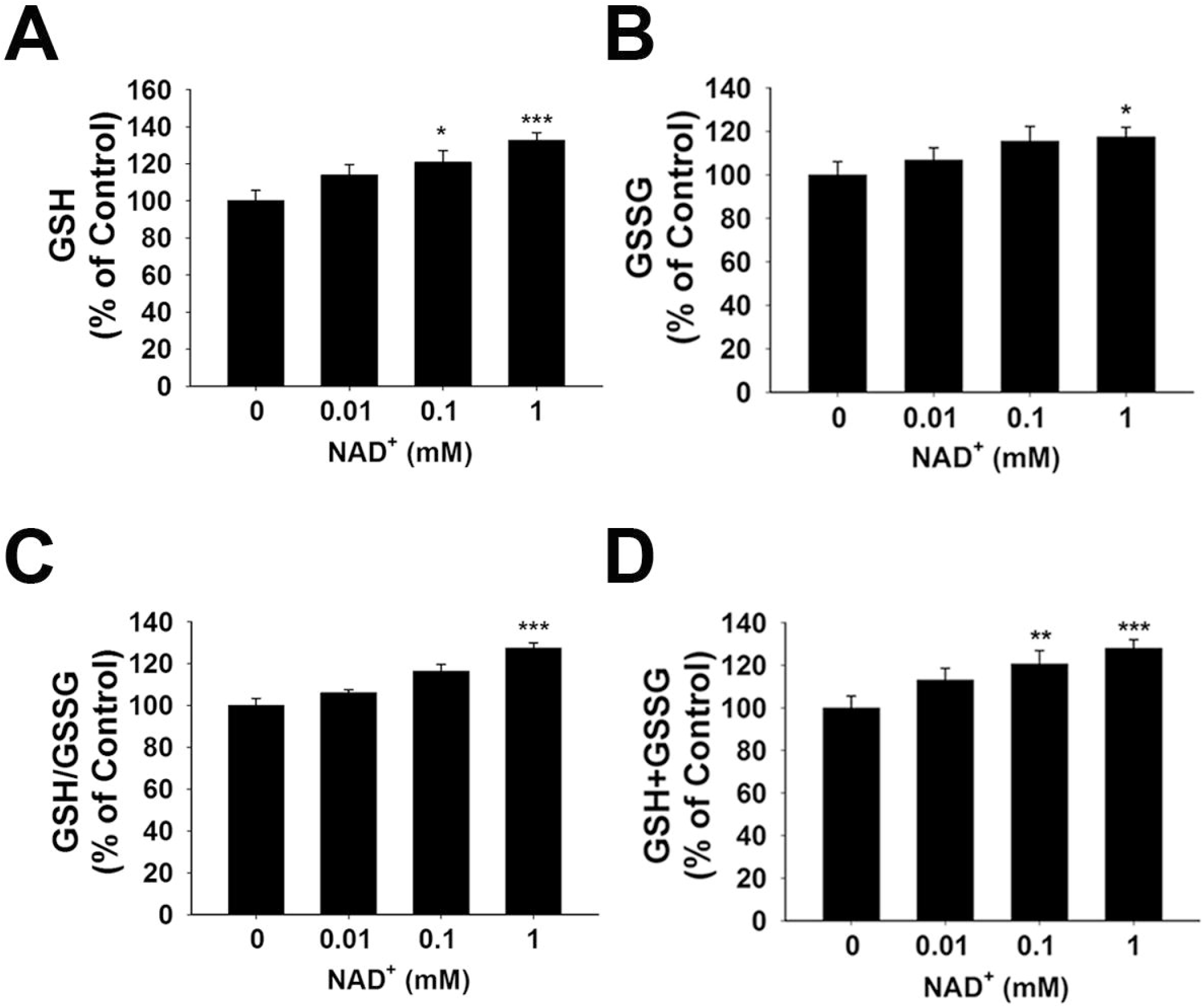
NAD^+^ treatment increased the glutathione levels and GSH/GSSG ratio in differentiated PC12 cells. (A) NAD^+^ treatment dose-dependently increased the GSH level in differentiated PC12 cells. (B) 1 mM NAD^+^ slightly increased the GSSG level in the cells, while lower concentrations of NAD^+^ did not affect the GSSG level. (C) 1 mM NAD^+^ significantly increased the GSH/GSSG ratio in the cells, while lower concentrations of NAD^+^ did not affect the GSH/GSSG ratio. (D) NAD^+^ treatment dose-dependently increased the total glutathione level in the cells. The assays on GSH and GSSG were conducted, after the cells were treated with NAD^+^ for 24 hours. N = 16. The data were pooled from four independent experiments. *, *p* < 0.05; **, *p* < 0.01; ***, *p* < 0.001.

### NAD^+^ treatment enhances the GSH/GSSG ratio and total glutathione levels by increasing both GCL levels and Nrf2 activity of PC12 cells

To investigate the mechanisms underlying the effects of NAD^+^ on the GSH/GSSG ratios, we determined the effects of NAD^+^ treatment on the intracellular NAD^+^ levels of both control and rotenone-treated cells. We found that treatment of the cells with NAD^+^ significantly increased the intracellular NAD^+^ levels in both control and rotenone-treated cells (Figure S1).

GCL is the key enzyme for GSH synthesis (Dickinson and Forman, 2002), while GSSG is originated from consumption of GSH in various antioxidant processes (Stryer, 1995). Our finding that NAD^+^ treatment can increase the total glutathione levels has suggested that the GCL activity may be increased in the NAD^+^-treated cells. GCL has two major subunits - GCLc and GCLm, both of which are important for GCL activity (Meister, 1991; Huang et al., 1993). Our real-time PCR assay showed that NAD^+^ significantly increased the mRNA levels of both GCLc and GCLm (Figures 2A, B). Our Western blot assays also showed that NAD^+^ significantly increased the protein levels of both GCLc and GCLm (Figures 2C-E).

**Figure 2.**
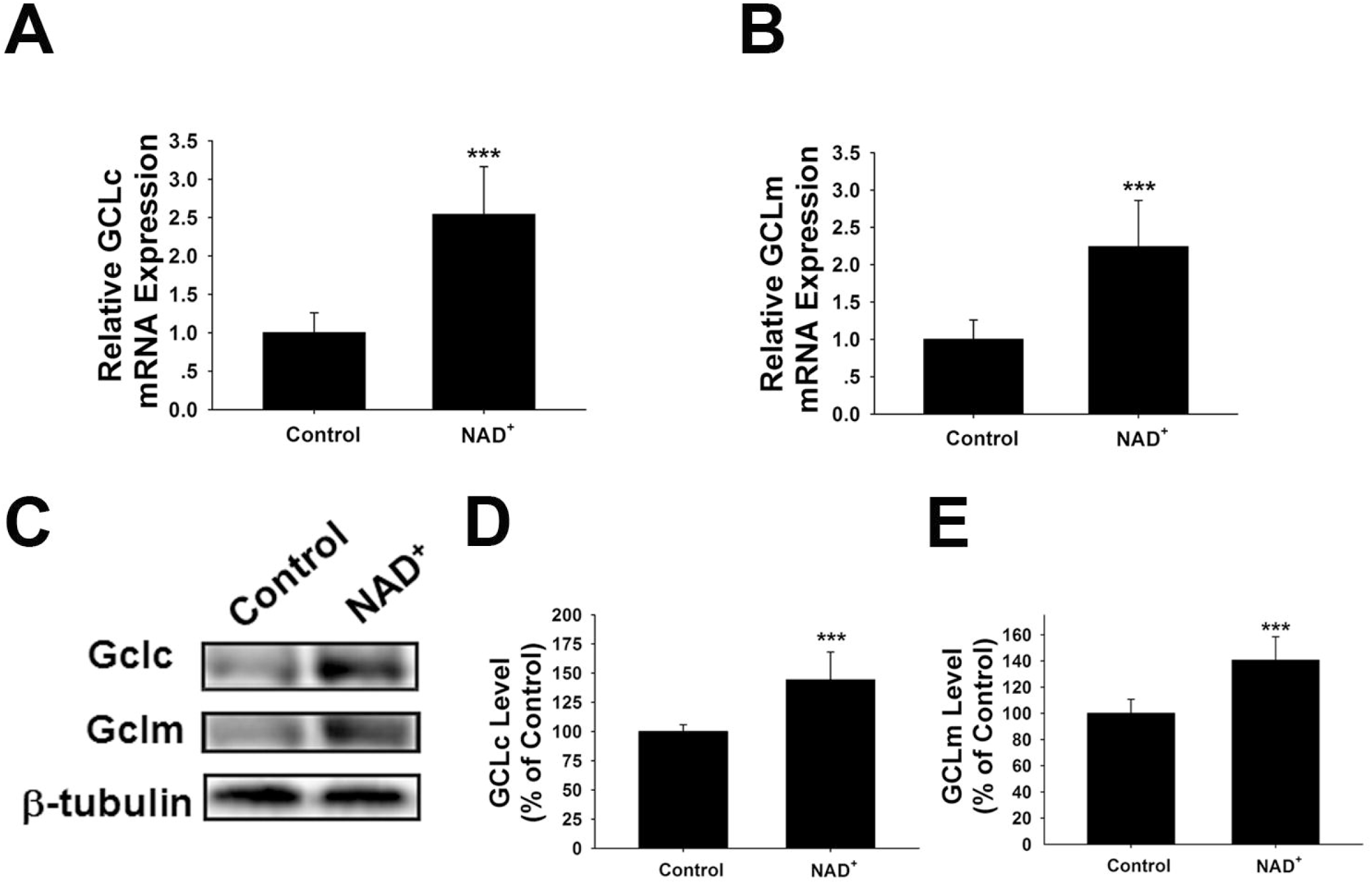
NAD^+^ treatment increased the gene expression and protein levels of GCLc and GCLm in PC12 cells. (A) Real-time PCR assays showed that NAD^+^ treatment led to a significant increase in the GCLc mRNA level, assessed at 6 hours after the NAD^+^ treatment. (B) Real-time PCR assays showed that NAD^+^ treatment led to significant increases in the GCLm mRNA levels, assessed at 6 hours after the NAD^+^ treatment. (C) Western blot assays showed that NAD^+^ treatment led to increased protein levels of GCLc and GCLm in PC12 cells, assessed at 12 hours after the NAD^+^ treatment. (D) Quantifications of the Western blots showed that NAD^+^ treatment led to a significant increase in the protein levels of GCLc. (E) Quantifications of the Western blots showed that NAD^+^ treatment led to a significant increase in the protein level of GCLm. N = 16. The data were pooled from four independent experiments. ***, *p* < 0.001.

Because induction of GCL expression have been reported to be dependent on Nrf2 (Chan and Kwong, 2000; Suh et al., 2004), we also determined the effects of NAD^+^ treatment on the Nrf2 mRNA levels and the nuclear translocation of Nrf2. Our study showed that NAD^+^ significantly increased not only the Nrf2 mRNA levels, but also the nuclear Nrf2 levels of the cells (Figures 3A-C).

**Figure 3.**
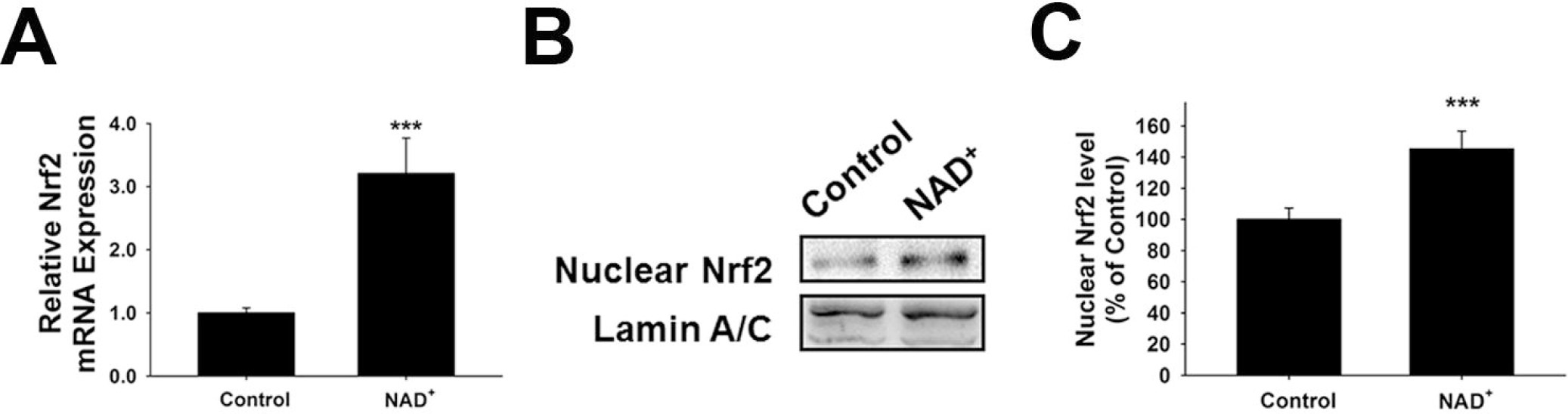
NAD^+^ treatment led to a significant increase in the mRNA levels and nuclear levels of Nrf2 of PC12 cells. (A) Real-time PCR assays showed that NAD^+^ treatment led to a significant increase in the Nrf2 mRNA level, assessed at 6 hours after the NAD^+^ treatment. (B) Western blot assays showed that NAD^+^ treatment led to an increased level of nuclear Nrf2 in the cells, assessed at 1 hour after the NAD^+^ treatment. (C) Quantifications of the Western blots showed that NAD^+^ treatment led to a significant increase in the nuclear levels of Nrf2. N = 12. *\*\*\*, p* < 0.01.

### SIRT2 mediates the NAD^+^-induced increase in GSH/GSSG ratio by regulating the Nrf2 and GCL in PC12 cells

It has been reported that NAD^+^-dependent enzymes SIRT1 and SIRT2 can affect the gene expression of antioxidant enzymes such as mitochondrial superoxide dismutase (SOD2) by affecting the transcriptional factors of FOXO family (Wang et al., 2007; Liu et al., 2013). Therefore, we investigated the roles of SIRT1 and SIRT2 in the NAD^+^-produced increase in the GSH/GSSG ratio. Our study did not observe any significant effects of the SIRT1 inhibitor Ex 527 (Selleck Chemicals, Houston, TX, USA) on the NAD^+^-produced increase in the GSH/GSSG ratio and the levels of GSH, GSSG; and total glutathione (Figure S2), thus arguing against the possibility that SIRT1 plays a major role in the NAD^+^-induced increases in the antioxidant capacity. In contrast, we found that SIRT2 inhibitor AGK2 was capable of abolishing the effects of NAD^+^ on the GSH/GSSG ratio and the levels of GSH, GSSG; and total glutathione of the cells (Figures 4A-D). AGK2 was also shown to block the NAD^+^-induced increases in both mRNA and protein levels of GCLc and GCLm (Figures 4E-I). Moreover, AGK2 treatment attenuated the NAD^+^-induced increases in the Nrf2 mRNA levels and the nuclear Nrf2 levels (Figures 4J-L).

**Figure 4.**
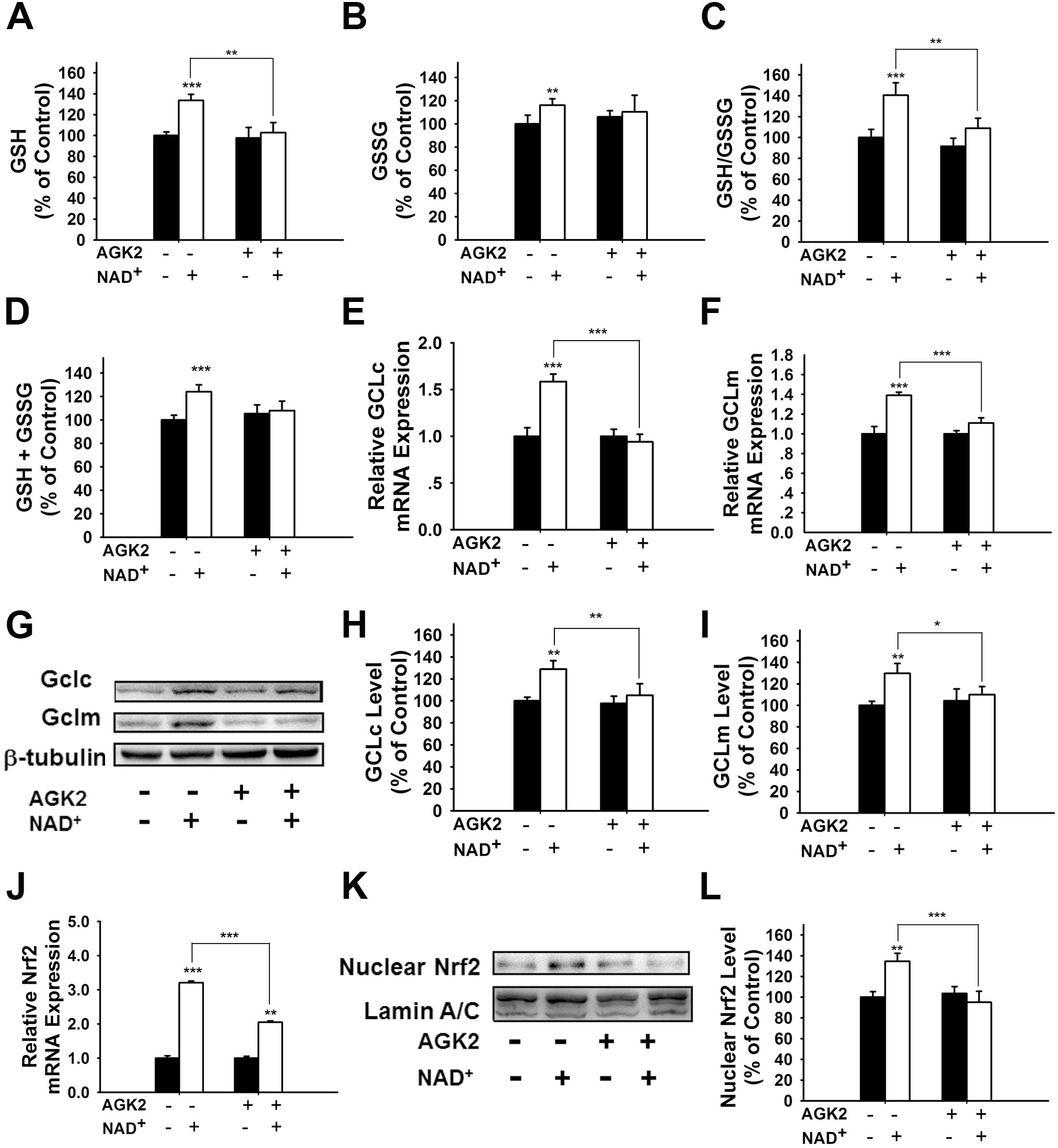
The SIRT2 inhibitor AGK2 prevented NAD^+^-induced increases in the glutathione levels, GSH/GSSG ratio, GCL level and nuclear Nrf2 level of PC12 cells. (A) AGK2 prevented the NAD^+^-induced increase in the GSH levels. (B) AGK2 prevented the NAD^+^-induced increase in the GSSG levels. (C) AGK2 prevented the NAD^+^-induced increase in the GSH/GSSG ratio. (D) AGK2 prevented the NAD^+^-induced increase in the total glutathione level. For A-D, the assays were conducted after the cells were treated with 1 mM NAD^+^ and 5 μM AGK2 for 24 hours. (E) AGK2 prevented the NAD^+^-induced increase in the GCLc mRNA level of the cells, assessed at 6 hours after the NAD^+^ treatment. (F) AGK2 prevented the NAD^+^-induced increase in the GCLm mRNA level of the cells, assessed at 6 hours after the NAD^+^ treatment. (G-I) AGK2 prevented the NAD^+^-induced increases in the protein levels of GCLc and GCLm in PC12 cells, assessed at 12 hours after the NAD^+^ treatment. (J) AGK2 prevented the NAD^+^-induced increase in the Nrf2 mRNA level of the cells, assessed at 6 hours after the NAD^+^ treatment. (K-L) AGK2 prevented the NAD^+^-induced nuclear translocation of Nrf2, assessed at 1 hour after the NAD^+^ treatment. The cells were treated with 1 mM NAD^+^ and 5 μM AGK2. N =16. The data were pooled from four independent experiments. *\*, p* < 0.05; *\*\*, p* < 0.01; *\*\*\*, p* < 0.001.

We further applied SIRT2 siRNA to investigate the role of SIRT2 in the NAD^+^-induced increases in glutathione synthesis. SIRT2 siRNA significantly decreased the expression of SIRT2 (Figure 5A). SIRT2 siRNA was also capable of preventing the NAD^+^-induced increases in the GSH/GSSG ratio and the levels of GSH, GSSG; and total glutathione (Figures 5B-E). We further applied real-time PCR assays to determine the effects of SIRT2 siRNA on the NAD^+^-induced increases in the mRNA levels of GCLc, GCLm and Nrf2, showing that SIRT2 silencing significantly decreased the NAD^+^-induced elevations of the mRNA levels of GCLc, GCLm and Nrf2 (Figures 5F-H).

**Figure 5.**
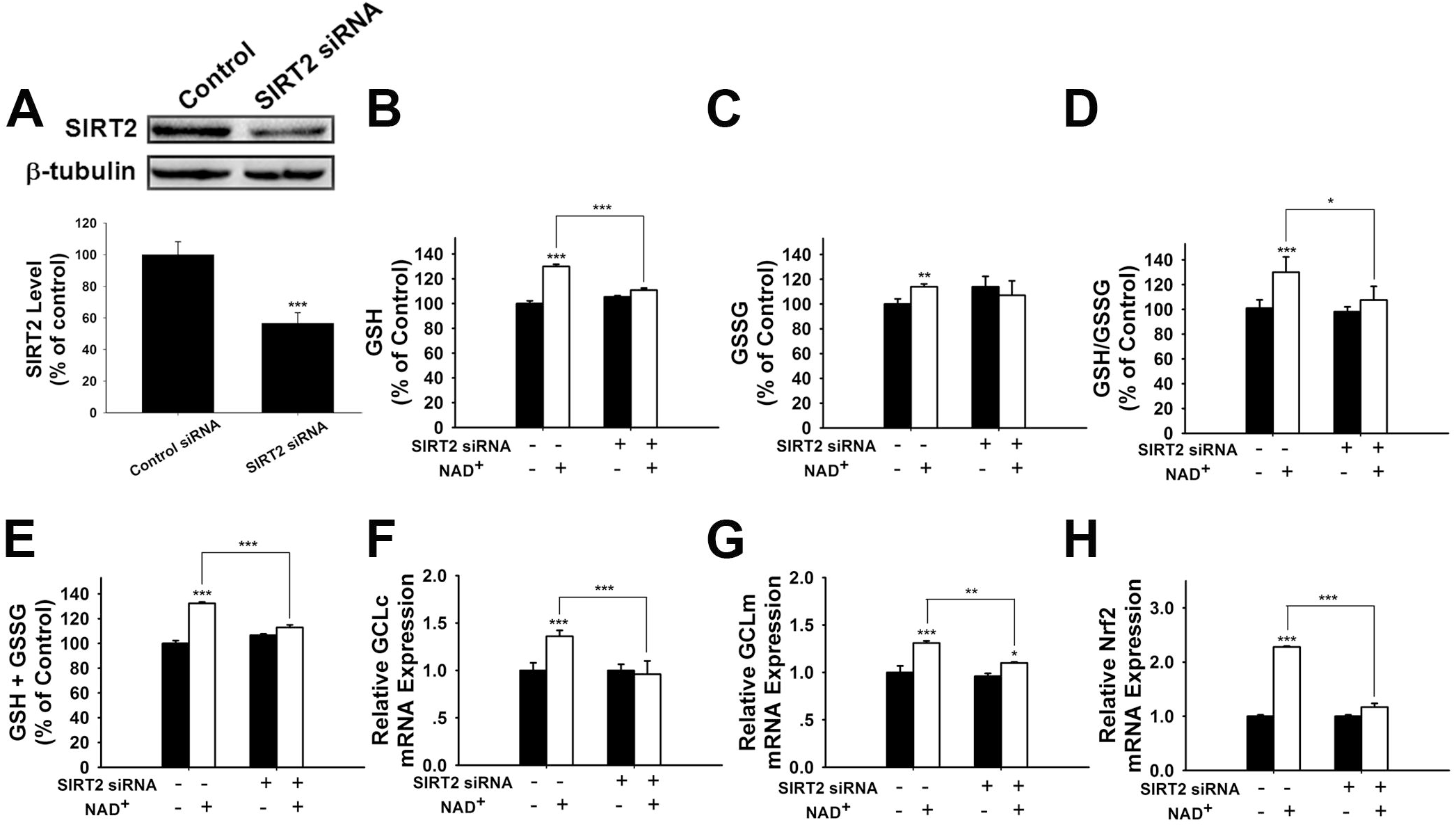
SIRT2 silencing prevented NAD^+^-induced increases in the glutathione levels, GSH/GSSG ratio, GCL level, and Nrf2 level of PC12 cells. (A) SIRT2 siRNA decreased the protein levels of SIRT2 in PC12 cells. (B) SIRT2 silencing prevented the NAD^+^-induced increase in the GSH level of the cells. (C) SIRT2 silencing prevented the NAD^+^-induced increase in the GSSG level of the cells. (D) SIRT2 silencing prevented the NAD^+^-induced increase in the GSH/GSSG ratio of the cells. (E) SIRT2 silencing prevented the NAD^+^-induced increase in the total glutathione level of the cells. (F) SIRT2 silencing prevented the NAD^+^-induced increase in the GCLc mRNA level of the cells. (G) SIRT2 silencing prevented the NAD^+^-induced increase in the GCLm mRNA level of the cells. (H) SIRT2 silencing prevented the NAD^+^-induced increase in the Nrf2 mRNA level of the cells. The cells were treated with SIRT2 siRNA for 6 hours, then the media was replaced with DMEM containing 10% fetal bovine serum. Then the cells were treated with or without NAD^+^. RT-PCR assays were carried out at 6 hours after the NAD^+^ treatment, while glutathione assays were carried out at 24 hours after the NAD^+^ treatment. N = 16. The data were pooled from four independent experiments. *\*, p* < 0.05; *\*\*, p* < 0.01; *\*\*\*, p* < 0.001.

### ERK activation plays a crucial role in the NAD^+^-induced increases in the GSH/GSSG ratio and Nrf2 activity of PC12 cells

It has been reported that pERK can induce Nrf2 activation by promoting its phosphorylation (Zipper and Mulcahy, 2000; Yang et al., 2011). Therefore, we determined if ERK phosphorylation plays a role in the NAD^+^-induced increase in the GSH/GSSG ratio. Our study showed that U0126, an inhibitor of the ERK pathway, attenuated the NAD^+^-induced increases in the GSH/GSSG ratio and the levels of GSH, GSSG; and total glutathione (Figures 6A-D). U0126 also blocked the NAD^+^-induced increases in the mRNA levels of GCLc, GCLm and Nrf2 (Figures 6E-G). Our Western blot assay showed that U0126 blocked the NAD^+^-induced nuclear translocation of Nrf2 (Figure 6H).

**Figure 6.**
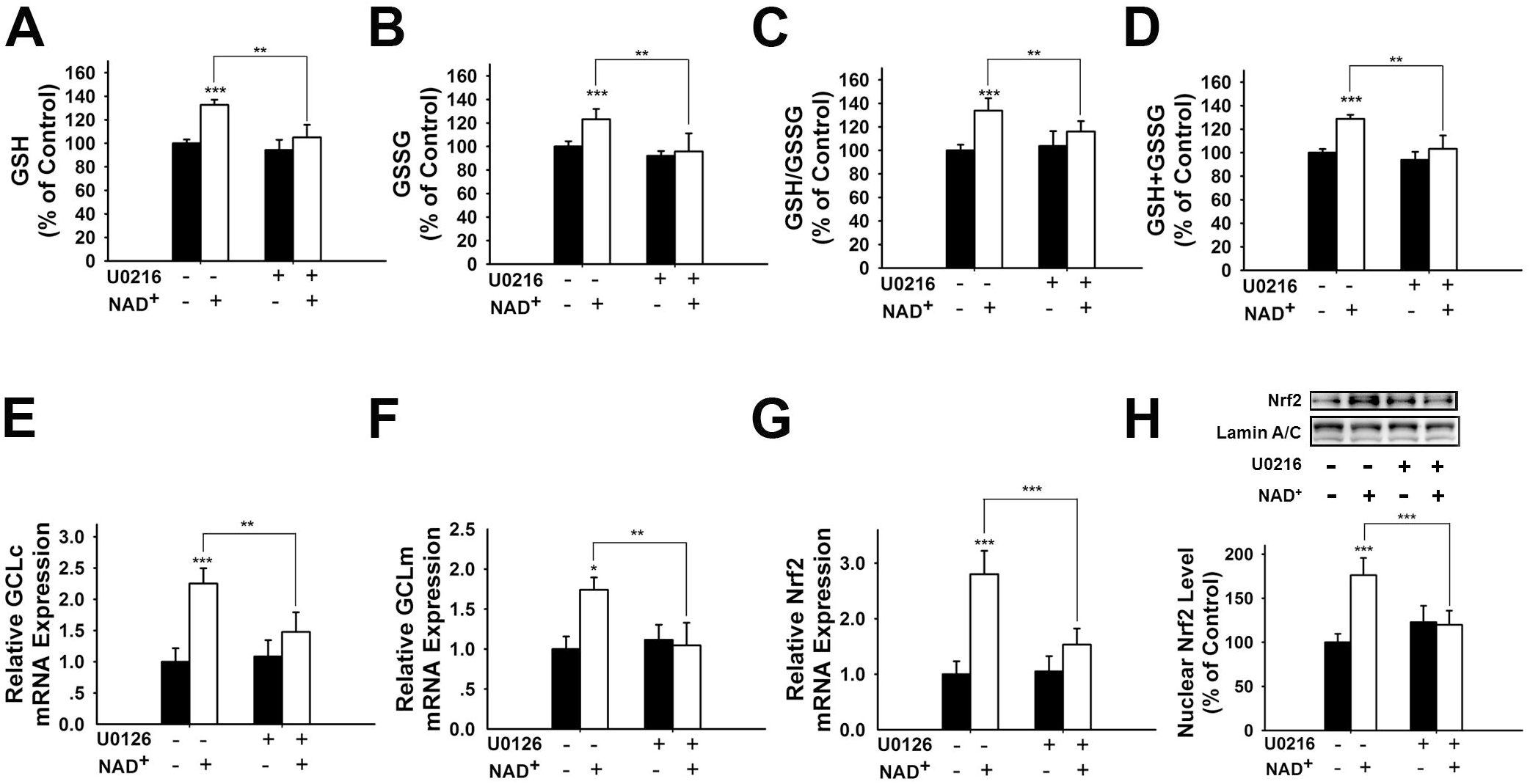
Inhibition of ERK signaling prevented NAD^+^-induced increases in the glutathione levels, GSH/GSSG ratio, GCL level, and Nrf2 levels of PC12 cells. (A) The MEK/ERK pathway inhibitor U0126 prevented the NAD^+^-induced increase in the GSH level of the cells. (B) U0126 prevented the NAD^+^-induced increase in the GSSG level of the cells. (C) U0126 prevented the NAD^+^-induced increase in the GSH/GSSG ratio of the cells. (D) U0126 prevented the NAD^+^-induced increase in the total glutathione level of the cells. (E) U0126 prevented the NAD^+^-induced increase in the GCLc mRNA level of the cells. (F) U0126 prevented the NAD^+^-induced increase in the GCLm mRNA level of the cells. (G) U0126 prevented the NAD^+^-induced increase in the Nrf2 mRNA level of the cells. (H) U0126 prevented the NAD^+^-induced increase in the nuclear Nrf2 protein level of the cells. The cells were treated with 1 mM NAD^+^ and 20 μM U0126. Glutathione assays were conducted at 24 hours after the NAD^+^ treatment. RT-PCR assays were conducted at 6 hours after the NAD^+^ treatment. Western blot assays were conducted at 1 hour after the NAD^+^ treatment. N = 16. The data were pooled from four independent experiments. *, *p* < 0.05; **, *p* < 0.01; ***, *p* < 0.001.

We also found that NAD^+^ treatment induced a significant increase in the pERK protein level in PC12 cells, which was attenuated by either SIRT2 siRNA or AGK2 (Figures 7A, B), suggesting that ERK was a downstream target of SIRT2 in the NAD^+^-induced increase in the GSH/GSSG ratio.

**Figure 7.**
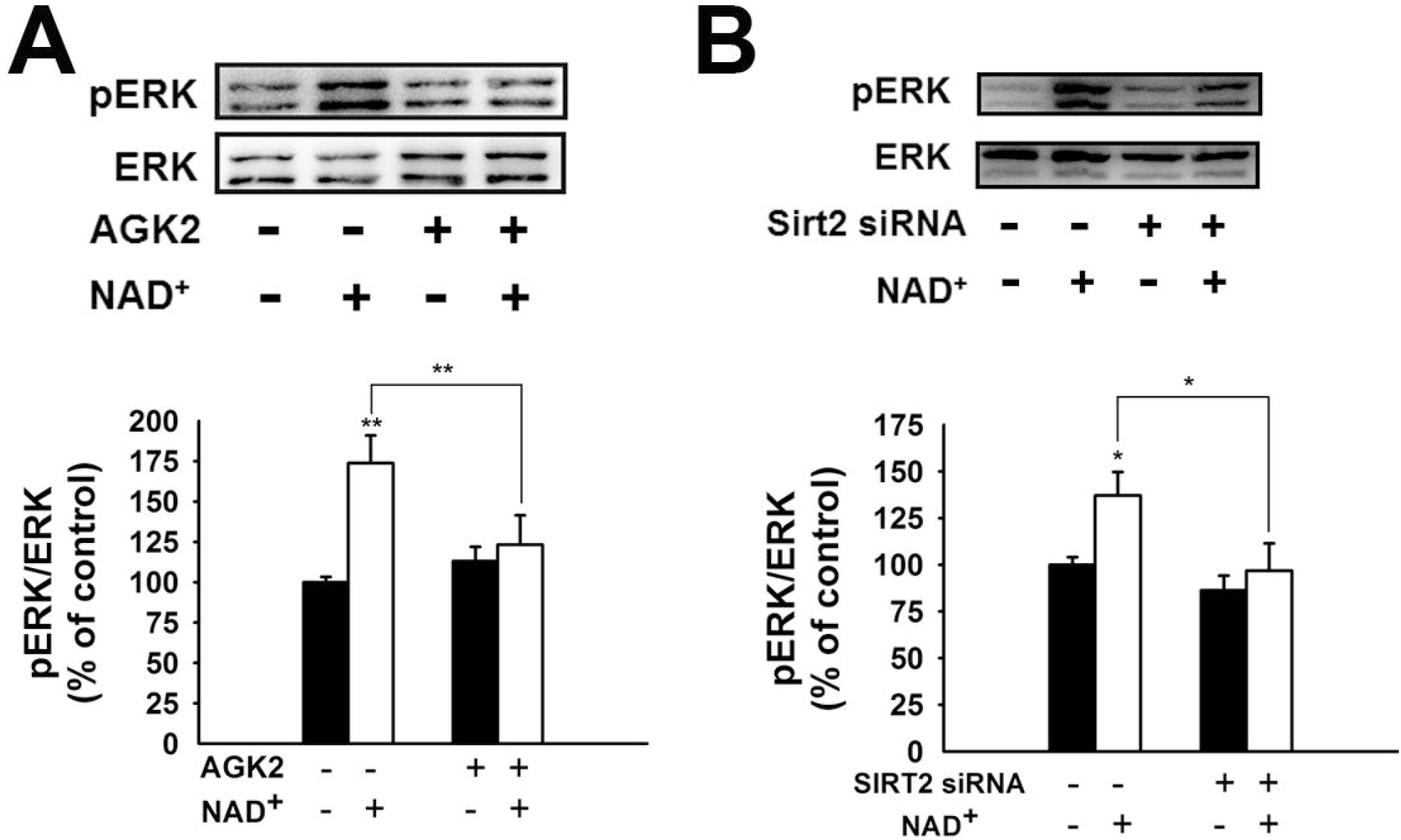
Both the SIRT2 inhibitor AGK2 and SIRT2 silencing prevented the NAD^+^-induced increase in phosphorylated ERK in PC12 cells. (A) AGK2 prevented the NAD^+^-induced increase in phosphorylated ERK. The cells were treated with 5 μM AGK2 and 1 mM NAD^+^ for 0.5 hour. (B) SIRT2 silencing prevented the NAD^+^-induced increase in phosphorylated ERK. After the cells were treated with SIRT2 siRNA for 6 hours, the media was replaced with DMEM containing 10% fetal bovine serum. Then the cells were treated with 1 mM NAD^+^ for 0.5 hour. *\*, p* < 0.05; *\*\*, p* < 0.01.

### NAD^+^ promoted the GSH synthesis in an adenosine-independent way

Our latest study reported that NAD^+^ induced increases in intracellular ATP levels under basal conditions through its degradation into adenosine (Zhang et al., 2018). Therefore, we investigated the role of adenosine in the NAD^+^-induced increases in the GSH synthesis and GSH/GSSG ratio. We found that NAD^+^ significantly increased the extracellular adenosine level, but only slight increased the intracellular adenosine levels (Figure S3). Consistent with the findings of Ramkumar et al. (Ramkumar et al., 1995), our study showed that adenosine increased the GSH levels and GSH/GSSG ratios (Figure S4 A-D).

Our current study has indicated that SIRT2 mediates the NAD^+^-induced increases in GSH synthesis. However, the mechanism underlying the adenosine-produced changes of GSH metabolism appears to be different from that underlying the NAD^+^-produced changes of GSH metabolism: The SIRT2 inhibitor AGK2 could not block the adenosine-produced changes of the glutathione metabolism (Figure S4 E-H).

## Discussion

The major findings of our current study include: First, NAD^+^ treatment can increase the GSH/GSSG ratio of PC12 cells under basal conditions, suggesting that NAD^+^ treatment can increase directly the antioxidant capacity of the cells; second, NAD^+^ treatment can increase both the mRNA and protein levels of GCL; third, NAD^+^ treatment can increase both the Nrf2 mRNA level and nuclear Nrf2 levels; fourth, SIRT2 mediates the NAD^+^-induced increase in the GSH/GSSG ratio, GCL levels and Nrf2 activity; fifth, ERK mediates the increases in the GSH/GSSG ratio and Nrf2 activity; and sixth, ERK is a downstream target of SIRT2 in the NAD^+^-produced changes of the glutathione metabolism. Collectively, our study has indicated that NAD^+^ can enhance directly the antioxidant capacity of the cells, which is mediated by SIRT2, ERK, and Nrf2.

Oxidative stress is a major pathological factor in numerous diseases and aging (Halliwell, 2005; Lin and Beal, 2006). Previous studies have suggested that NAD^+^ treatment can decrease oxidative cell death indirectly by such mechanisms as enhancement of SIRT1 activity and prevention of glycolytic inhibition, mitochondrial permeability transition, mitochondrial depolarization and nuclear translocation of apoptosis-inducing factor (AIF) (Alano et al., 2004; Alano et al., 2010; Hong et al., 2014). However, it remains unclear if NAD^+^ may attenuate oxidative damage by increasing directly antioxidant capacity of normal cells. Our current study has provided the first evidence suggesting that NAD^+^ treatment can enhance directly the antioxidant capacity of cells under basal conditions: NAD^+^ treatment can significantly increase the GSH/GSSG ratio of the cells. Our findings have suggested that NAD^+^ administration may produce its profound protective effects by increasing directly the antioxidant capacity of normal cells in the models of various diseases and aging, which can lead to increased resistance of the normal cells to oxidative insults.

Our study has indicated that NAD^+^ can significantly increase the mRNA levels and the protein levels of both GCLc and GCLm of the cells. These observations have suggested that NAD^+^ treatment leads to increased GSH synthesis by enhancing the mRNA levels and the protein levels of GCL. Previous studies have suggested that Nrf2 is a transcriptional factor for the gene expression of GCL (Chan and Kwong, 2000; Suh et al., 2004). Our current study has shown that NAD^+^ can increase both Nrf2 mRNA level and the nuclear Nrf2 levels, which have suggested that NAD^+^ treatment leads to increased GCL levels at least partially by activating Nrf2.

Our current study has also indicated that SIRT2, rather than SIRT1, mediates the NAD^+^-induced increase in the antioxidant capacity: Both SIRT2 silencing and the SIRT2 inhibitor AGK2 can virtually abolish the effects of NAD^+^ on the GSH/GSSG ratio, the GCL levels and the Nrf2 activity. This finding, together with the previous studies indicating the capacity of SIRT2 to modulate the gene expression of SOD2 by affecting the activity of FOXO3a (Wang et al., 2007; Liu et al., 2013), has suggested that SIRT2 is an important modulator of antioxidant capacity of cells.

SIRT2 is a NAD^+^-dependent enzyme (Schemies et al., 2010). A recent study has shown that decreased NAD^+^ levels can produce decreased SIRT2 activity (Skoge et al., 2014). Therefore, in our current study, the NAD^+^ treatment-induced increases in the intracellular NAD^+^ levels could lead to increased SIRT2 activity, resulting in the increases in the Nrf2 activity, GCL levels and GSH/GSSG ratios. It is seemingly puzzling that SIRT2, but not SIRT1, mediates the NAD^+^-induced changes of the glutathione metabolism. We propose the following mechanisms for accounting for this finding: SIRT2 is a cytosolic enzyme, while SIRT1 is mainly a nuclear enzyme (Houtkooper et al., 2012). NAD^+^ treatment may selectively increase cytosolic NAD^+^ levels, leading to a selective increase in SIRT2 activity. Future studies are warranted to further elucidate the mechanisms underlying this finding.

Our study has indicated that ERK links SIRT2 and the NAD^+^ treatment-produced changes of the GSH/GSSG ratio, the GCL levels and the Nrf2 activity in our experimental model: First, both SIRT2 siRNA and AGK2 can prevent the NAD^+^-induced increase in ERK phosphorylation, indicating that SIRT2 mediates the NAD^+^-induced ERK phosphorylation; and second, ERK inhibition can also significantly attenuate the NAD^+^-induced increases in the GSH/GSSG ratio and the Nrf2 activity. These findings are consistent with previous reports regarding the linkage between SIRT2 and ERK (Koh, 2012; Sung et al., 2012; Yeung et al., 2015; Eo et al., 2016; Jeong and Cho, 2017). Collectively, our study has suggested the following mechanism underlying the effects of NAD^+^ treatment on the glutathione metabolism (Figure 8): NAD^+^ treatment produces increased SIRT2 activity that causes ERK activation, leading to increased Nrf2 activity, resulting in increased levels of GCL and glutathione synthesis. Future studies are warranted to determine if this pathway occurs *in vivo*.

**Figure 8.**
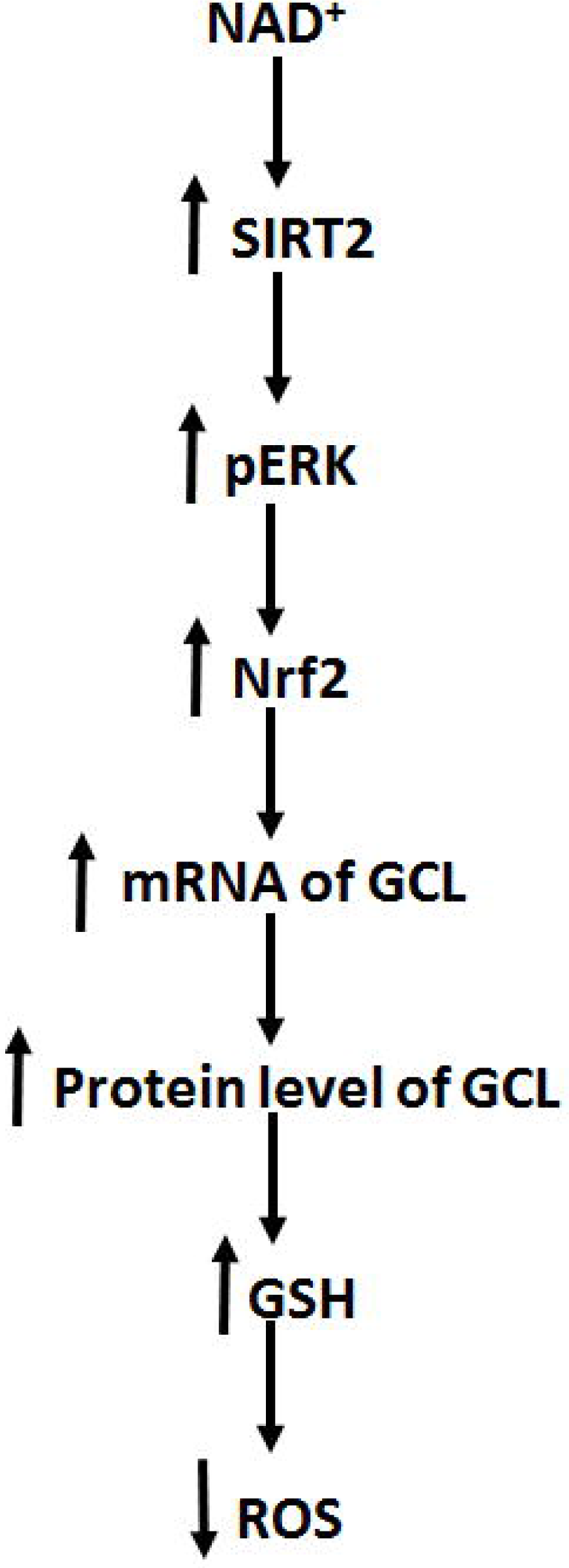
Diagrammatic presentation of the mechanisms underlying the relationships among NAD^+^, ERK, SIRT2, Nrf2, GCL, glutathione metabolism, and antioxidant capacity of cells.

Our recent study reported that NAD^+^ induced increases in intracellular ATP levels under basal conditions through its degradation into adenosine (Zhang et al., 2018). Our current study has suggested that NAD^+^ increases the GSH synthesis of the cells under basal conditions by an adenosine-independent pathway: While NAD^+^ and adenosine can increase GSH synthesis. AGK2 can reverse only the NAD^+^ treatment-produced changes of the glutathione metabolism, but not those induced by adenosine treatment. Moreover, the concentrations of adenosine generated from the degradation of 1 mM NAD^+^ are much lower than 0.01 mM, which is not sufficient to increase GSH synthesis.

Our previous study has suggested that NADH can significantly increase the levels of GSH and GSSG of PC12 cells (Cao et al., 2016). However, it is noteworthy that NAD^+^, but not NADH, can increase GSH/GSSG ratio - a major index of antioxidant capacity of cells. These observations have suggested that NAD^+^ is superior over NADH at least in enhancing antioxidant capacity of normal cells, which may account for the following important observations regarding the biological effects of NAD^+^ and NADH: Compared with NADH, NAD^+^ has shown profound protective effects in much more models of diseases and aging (Ying, 2008). Moreover, our current study has indicated that the NAD^+^-induced increase in glutathione synthesis is Akt-independent (Figure S6), while the NADH-induced increase in glutathione synthesis is Akt-dependent (Cao et al., 2016).

## Acknowledgment

The authors would like to acknowledge the financial support by a Major Special Program Grant of Shanghai Municipality (Grant # 2017SHZDZX01) (to W.Y.), and a Major Research Grant from the Scientific Committee of Shanghai Municipality #16JC1400500 and #16JC1400502 (to W.Y.).

## Legends of Supplemental Figures

**Figure S1. NAD^+^ treatment led to a significant increase in the intracellular NAD^+^ level in both control and rotenone-treated PC12 cells.** The cells were treated with 1 mM NAD^+^ and 0.75 μM rotenone. One hour after the drug treatment, the intracellular NAD^+^ levels in the cells were determined by the cycling assays. N = 16. The data were pooled from four independent experiments. *\*\*\*, p* < 0.001.

**Figure S2. EX 527, a SIRT1 inhibitor, did not prevent the NAD^+^-induced increases in the GSH level, total glutathione level, and GSH/GSSG ratio in PC12 cells.** (A) EX 527 did not prevent the NAD^+^-induced increase in the GSH level of the cells. (B) EX 527 did not prevent the NAD^+^-induced increase in the GSSG level of the cells. (C) EX 527 did not prevent the NAD^+^-induced increase in the GSH/GSSG ratio of the cells. (D) EX 527 did not prevent the NAD^+^-induced increase in the total glutathione level of the cells. N = 16. The data were pooled from four independent experiments. *\*, p* < 0.05; *\*\*, p* < 0.01.

**Figure S3. NAD^+^ treatment led to a significant increase in the intracellular and extracellular adenosine levels in PC12 cells.** After the cells were treated with 1 mM NAD^+^ for 24 h, the intracellular and extracellular adenosine levels were determined. N = 16. The data were pooled from four independent experiments. *\*\*\*, p* < 0.001.

**Figure S4. AGK2 could not block the adenosine-produced increases in the glutathione level and GSH/GSSG ratio.** (A) Adenosine treatment increased the GSH levels in the cells. (B) Ten μM adenosine slightly decreased the GSSG levels while higher concentrations of adenosine did not affect the GSSG levels. (C) Adenosine significantly increased the GSH/GSSG ratio in the cells. (D) Adenosine treatment dose-dependently increased the total glutathione level of the cells. (E) AGK2 did not prevent the adenosine-induced increase in the GSH level of the cells. (F) The GSSG level was not affected by AGK2 or adenosine. (G) AGK2 did not prevent the adenosine-induced increase in the GSH/GSSG ratio of the cells. (H) AGK2 did not prevent the adenosine-induced increase in the total glutathione level of the cells. The cells were treated with 1 mM adenosine with or without 10 AGK2 for 24 hours. N = 12. The data were pooled from four independent experiments. *\*, p* < 0.05; *\*\*, p* < 0.01; *\*\*\*, p* < 0.001.

**Figure S5. LY294002, a PI3K/Akt pathway inhibitor, did not prevent the NAD^+^-induced increases in the glutathione levels and GSH/GSSG ratio in PC12 cells.** (A) LY294002 did not prevent the NAD^+^-induced increase in the GSH level of the cells. (B) LY294002 did not prevent the NAD^+^-induced increase in the GSSG level of the cells. (C) LY294002 did not prevent the NAD^+^-induced increase in the GSH/GSSG ratio of the cells. (D) LY294002 did not prevent the NAD^+^-induced increase in the total glutathione level of the cells. The assays were conducted, after the cells were treated with 1 μM LY294002 and 1 mM NAD^+^ for 24 hours. N = 16. The data were pooled from four independent experiments. *\*, p* < 0.05; *\*\*, p* < 0.01.

