## Supplementary figures and images for "SIRT2, ERK and Nrf2 mediate NAD+ treatment-induced increase in the antioxidant capacity of differentiated PC12 cells under basal conditions"

### Supplemental Figure 1

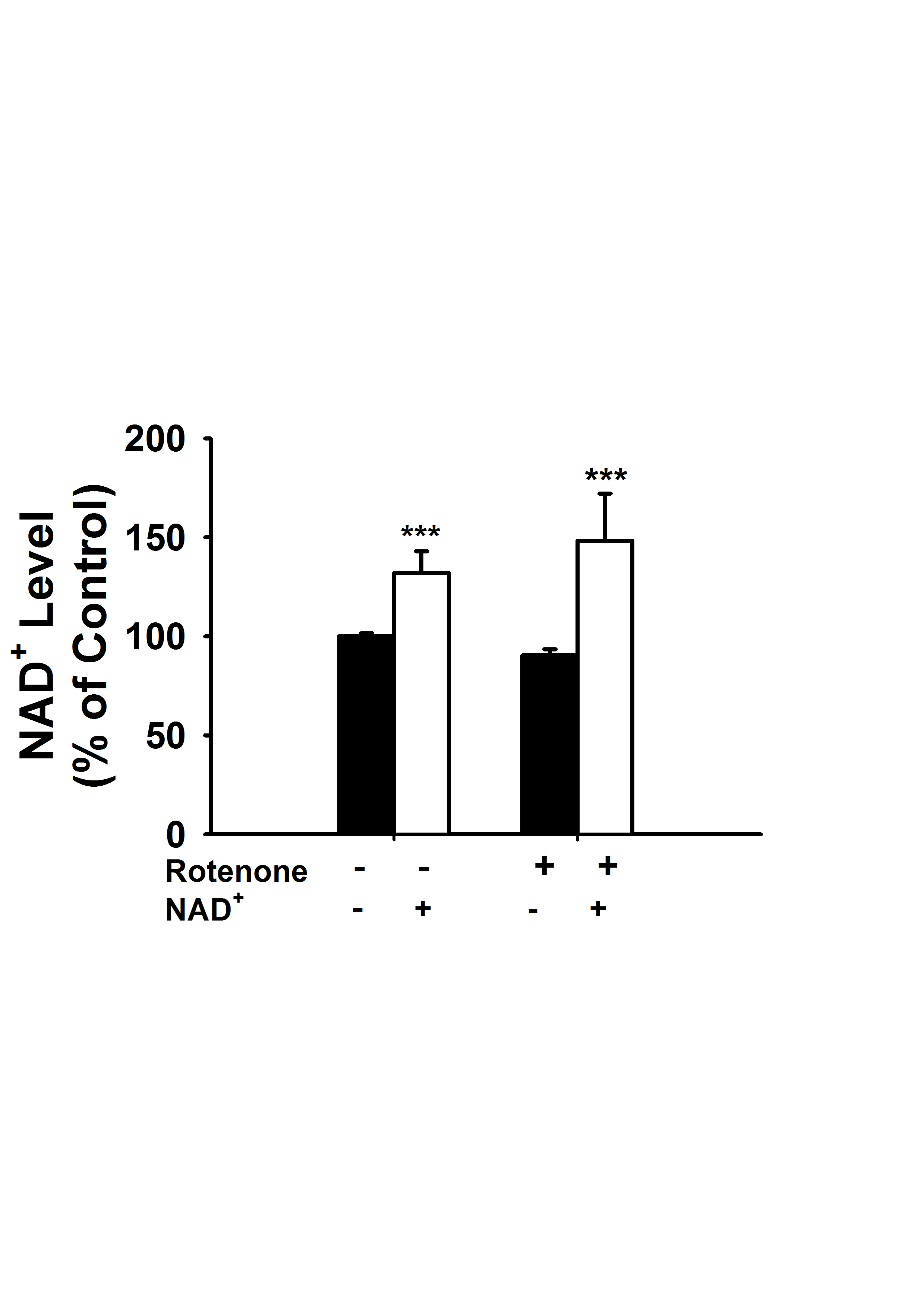

### Supplemental Figure 2

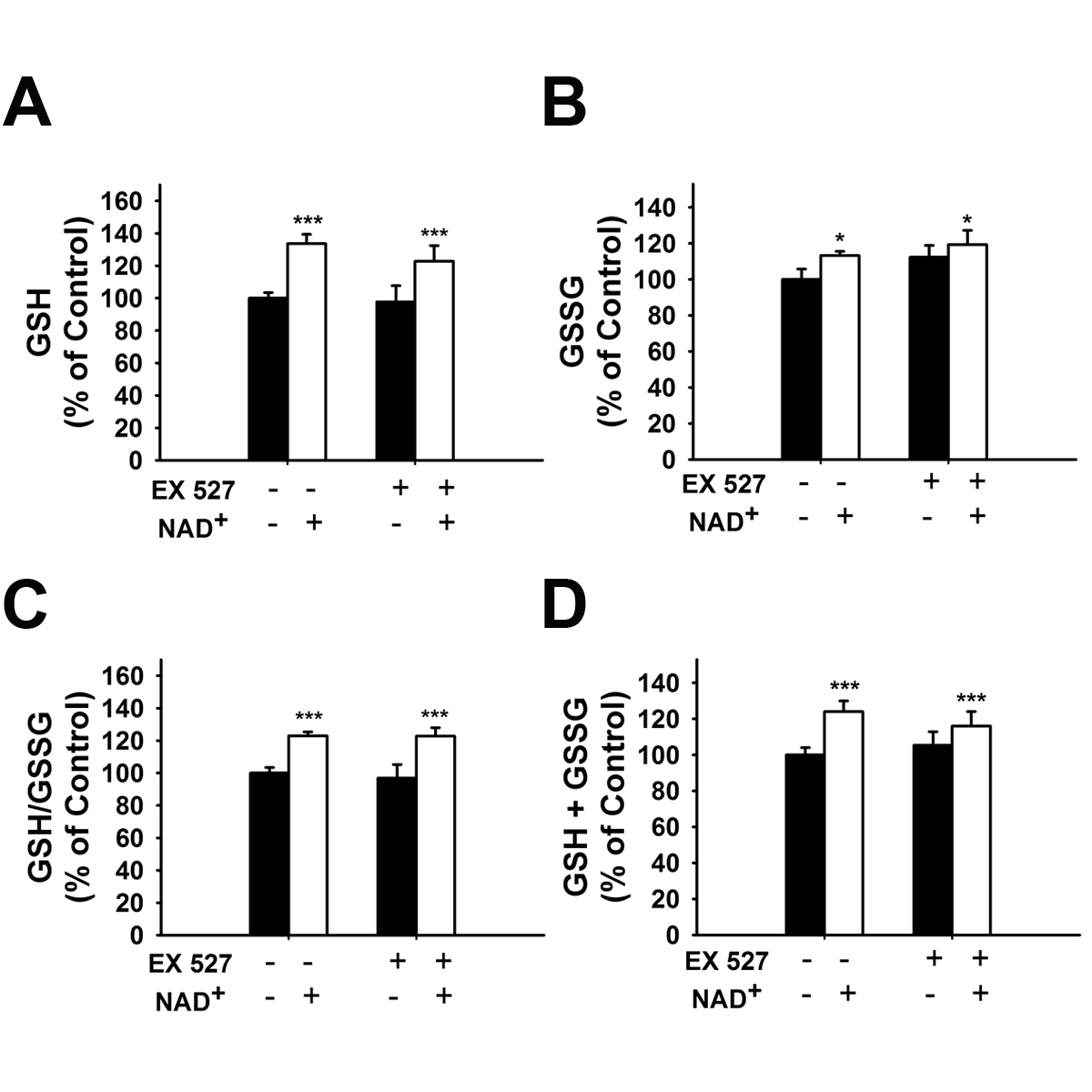

### Supplemental Figure 3

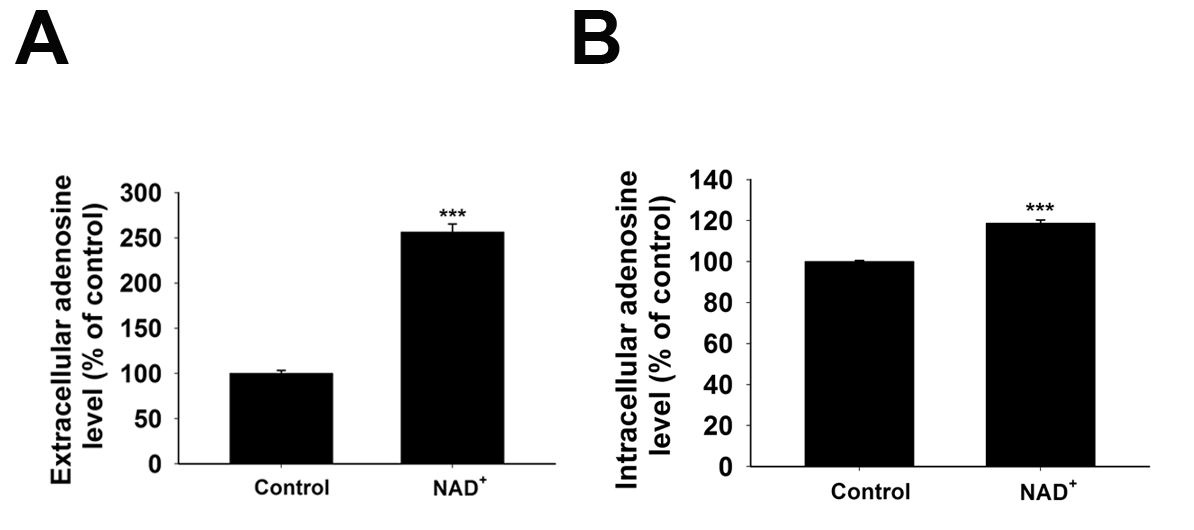

### Supplemental Figure 4

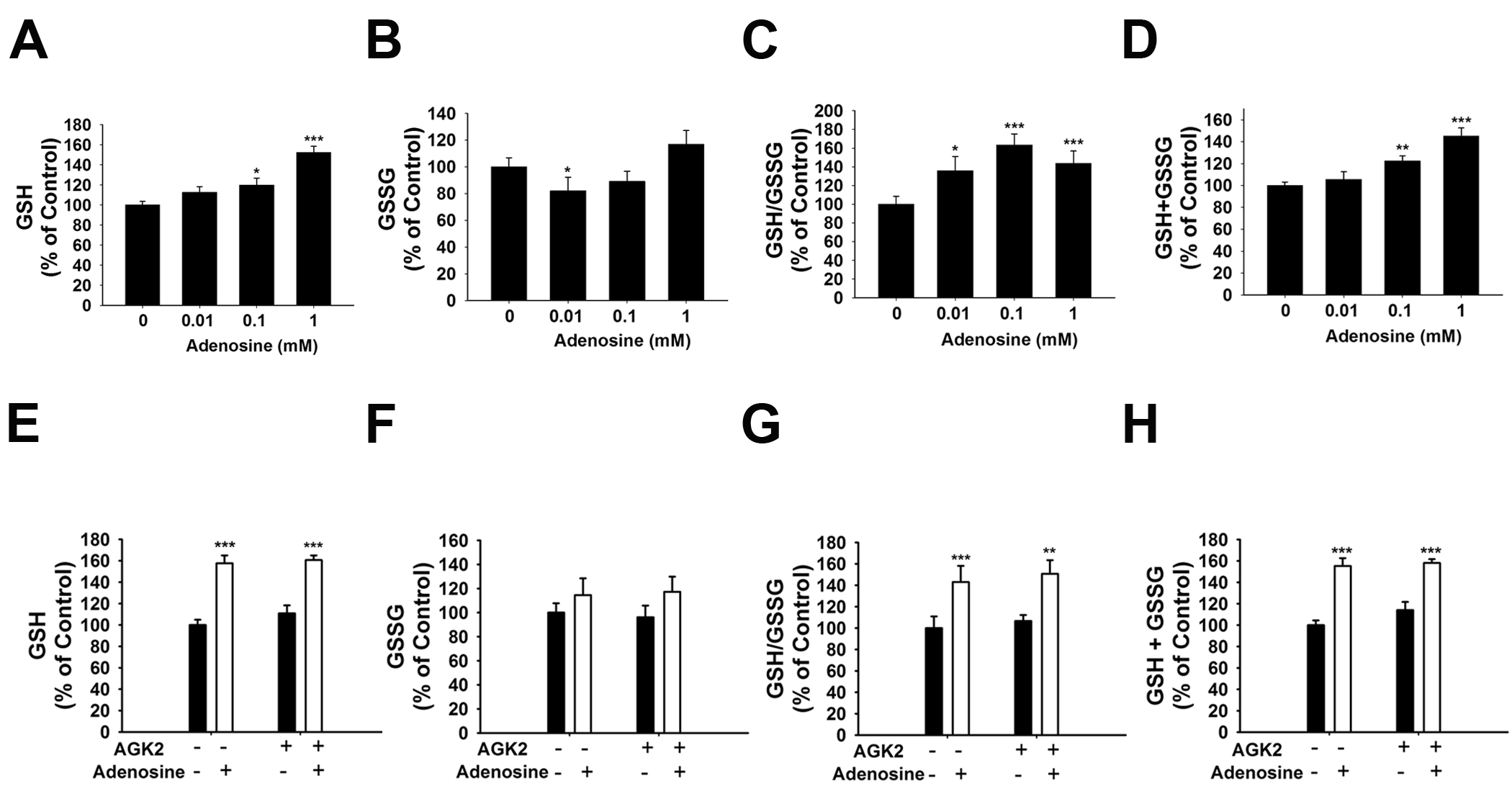

### Supplemental Figure 5

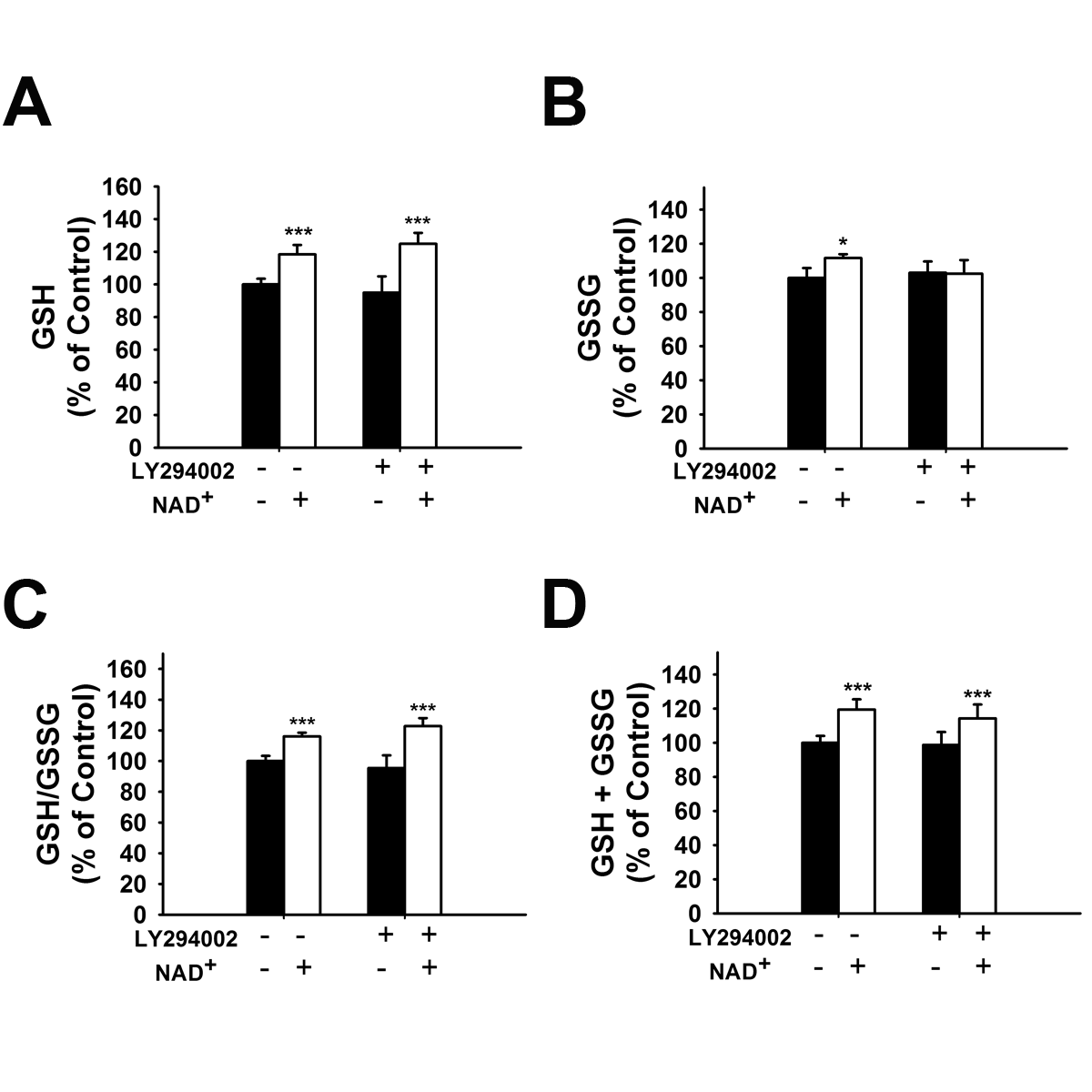
